# Circular RNA as a potential biomarker for forensic age prediction using multiple machine learning models: A preliminary study

**DOI:** 10.1101/2020.11.10.376418

**Authors:** Junyan Wang, Chunyan Wang, Lihong Fu, Qian Wang, Guangping Fu, Chaolong Lu, Jin Feng, Bin Cong, Shujin Li

## Abstract

In forensic science, accurate estimation of the age of a victim or suspect can facilitate the investigators to narrow a search and aid in solving a crime. Aging is a complex process associated with various molecular regulation on DNA or RNA levels. Recent studies have shown that circular RNAs (circRNAs) upregulate globally during aging in multiple organisms such as mice and elegans because of their ability to resist degradation by exoribonucleases. In the current study, we attempted to investigate circRNAs’ potential capability of age prediction. Here, we identified more than 40,000 circRNAs in the blood of thirteen Chinese unrelated healthy individuals with ages of 20-62 years according to their circRNA-seq profiles. Three methods were applied to select age-related circRNAs candidates including false discovery rate, lasso regression, and support vector machine. The analysis uncovered a strong bias for circRNA upregulation during aging in human blood. A total of 28 circRNAs were chosen for further validation in 50 healthy unrelated subjects aged between 19 and 72 years by RT-qPCR and finally, 7 age-related circRNAs were chosen for final age prediction models. Several different algorithms including multivariate linear regression (MLR), regression tree, bagging regression, random forest regression (RFR), and support vector regression (SVR) were compared based on root mean square error (RMSE) and mean average error (MAE) values. Among five modeling methods, random forest regression (RFR) performed better than the others with an RMSE value of 5.072 years and an MAE value of 4.065 years (R^2^ = 0.902). In this preliminary study, we firstly used circRNAs as additional novel age-related biomarkers for developing forensic age estimation models. We propose that the use of circRNAs to obtain additional clues for forensic investigations and serve as aging indicators for age prediction would become a promising field of interest.

**Author summary:** In forensic investigations, estimation of the age of biological evidence recovered from crime scenes can provide additional information such as chronological age or the appearance of a culprit, which could give valuable investigative leads especially when there is no eyewitness available. Hence, generating an accurate model for age prediction using body fluids such as blood commonly seen at a crime scene can be of vital importance. Various molecular changes on DNA or RNA levels were discovered that they upregulated or downregulated during a person’s lifetime. Although some biomarkers have been proved to be associated with aging and used to predict age, several disadvantages such as low sensitivity, prediction accuracy, instability and susceptibility of diseases or immune states, thus limiting their applicability in the field of age estimation. Here, we utilized a novel biomarker namely circular RNA (circRNA) to generate highly accurate age prediction models. We propose that circRNA is more suitable for forensic degradation samples because of its unique molecular structure. This preliminary research offers a new thought for exploring potential biomarker for age prediction.

## Introduction

In the forensic field, estimating the age of a recovered stain’s donor is extremely valuable because it can provide additional information such as the appearance of an unknown individual thus narrowing down the number of criminal suspects or missing person[1]. Traditional age estimation mostly relies on osteology and odontology methods which are restricted to a nearly complete skeleton and are influenced by subjective factors with low accuracy[2]. Additionally, in most cases, only fragmentary remains are left by the perpetrator after committing a crime which makes some biological evidence more important, especially blood which is one of the most common body fluids at crime scenes that deserved to be investigated in forensic fields.

To predict an individual’s age from DNA, considerable attempts have been made to find biomarkers as indicators of chronological age. Several molecular methods have been proposed such as telomere shortening[3, 4], mitochondrial DNA deletion[5, 6], signal-joint T-cell receptor excision circle (sjTRECs)[7], and DNA methylation[8–11]. Among these biomarkers, DNA methylation pattern was considered as the most promising age-predictive biomarkers for forensic and clinic use due to its high prediction accuracy. In particular, Zbiec-Piekarska et al[12]. conducted a research based on pyrosequencing data for two CpGs in the ELOVL2 gene achieved relatively high prediction accuracy with a mean absolute deviation (MAD) from chronological age of 5.03 years. Another model based on 5 CpG sites was reported by Zbiec-Piekarska, showing high prediction accuracy with a MAD of 3.9 years. Hence, several sites of gene ELOVL2 is believed to be the most hopeful locus for age prediction[13]. An age prediction model of 83 newly CpG sites identified through Epigenome-wide association analysis with a correlation of 0.99 and the error of 0.23 years indicated that the chronological age can be accurately predicted among children and adolescents using DNA methylation biomarker[14]. Although a DNA methylation-based age prediction method can be developed in a way relatively high accuracy for individual age estimation. It cannot be denied that DNA methylation change pattern can be greatly influenced by several factors such as smoking, nutrition, and diverse diseases giving rise to the inaccuracy of quantification results especially when the age increased, the bias bigger[15]. Furthermore, the problem of a critical level of degradation in the amount of full-length DNA after conventional bisulfite treatment is yet to be addressed[10]. In light of various restrictions and challenges mentioned above, looking for other appropriate biomarkers with higher prediction accuracy and stability in human blood is of great significance to forensic age estimation.

With the advent of RNA next-generation sequencing, circular RNAs (circRNAs) have emerged as an interesting RNA species[16, 17]. CircRNA is a class of newly discovered noncoding RNA (ncRNA) recent years, representing as a special covalent loop without a 5’cap or 3’ tail whose unique feature enhances their ability to resist the degradation of exonucleases, thus contributes to their stability compared to their mRNA counterparts[18, 19]. Apart from the property of resistance to RNase R digestion, circRNA also exhibits other biological characteristics like widespread expression, cell-specificity, tissue-specificity, and developmental stage-specific expression patterns[20].

Its feature of developmental stage-specific expression has been confirmed across various model organisms such as mice[21], flies[22], and elegans[23], indicating a potential role as biomarkers of chronological age. Hall et al.[24] identified 38 circRNAs that were differentially expressed between day 10 and 40 from RNA-seq profiles of Drosophila photoreceptor neurons. After detecting the global profiles of circRNAs in C. elegans from the fourth larval stage (L4) through 10-day old adults, Cortes-Lopez et al.[25] found a massive accumulation of circRNAs during aging. Although the fact that circRNAs are observed to accumulate during aging in various organisms and human senescent cells[26], to this day, however, research focusing on circRNAs for age prediction has not been carried out yet in the field of forensic.

In this regard, we analyzed circRNA-seq profiles from 13 blood samples of unrelated Chinese aged between 20 and 62 years, focusing on the potential links between chronological age and the expression of circular RNAs in human blood. The present study aims to build a novel, accurate, and reliable age prediction model based on a subset of age-related circRNAs. Notably, when it comes to constructing a prediction model, simple univariate or multiple linear methods are more likely to be doubtful of their ability to explain such a complicated connection between age and expression levels of molecular markers. However, machine learning has gained a place in medicine and captured the interest of medical researchers[27]. With the introduction of various machine-learning methods, the prediction accuracy of models has been significantly improved for age prediction, suggesting that machine learning algorithms can be more robust and efficient to obtain more accurate estimation models[28–30]. Thus, we built five different models including multivariate linear regression, regression tree, bagging regression, random forest regression (RFR), and support vector regression (SVR) based on seven highly age-associated circRNAs. To the best of our knowledge, this is the first study that uses circRNAs in blood as indicators together with machine learning methods to develop prediction models for forensic individual age estimation.

## Results

### Selection of circRNAs differentially expressed during aging

Peripheral whole blood from thirteen unrelated Chinese aged between 20 and 62 years was collected (S1 Table). A total of 45697 circRNAs were identified by circRNA high-throughput sequencing from comprehensive circRNA expression profiles among 13 unrelated Chinese with age ranging from 20 to 62 years (Fig 1a), demonstrating a high abundance of circRNA in human peripheral whole blood with an average of 26719 read counts for each individual (Fig 1b). This was in line with previous study conducted by Memczak et al. who discovered > 15-fold higher general circRNA expression in blood compared to the liver tissues and a level comparable to the circRNA rich cerebellum, indicating circRNAs could be used as biomarker molecules in standard clinical blood samples[31]. Additionally, circRNAs were distributed across various genomic regions, but most commonly from protein coding regions, where over 80% of circRNAs originated (Fig 1c). We also observed exons and introns accounting for a higher proportion (more than 90% in each sample) than intergenic regions (Fig 1d).

**Fig 1.**
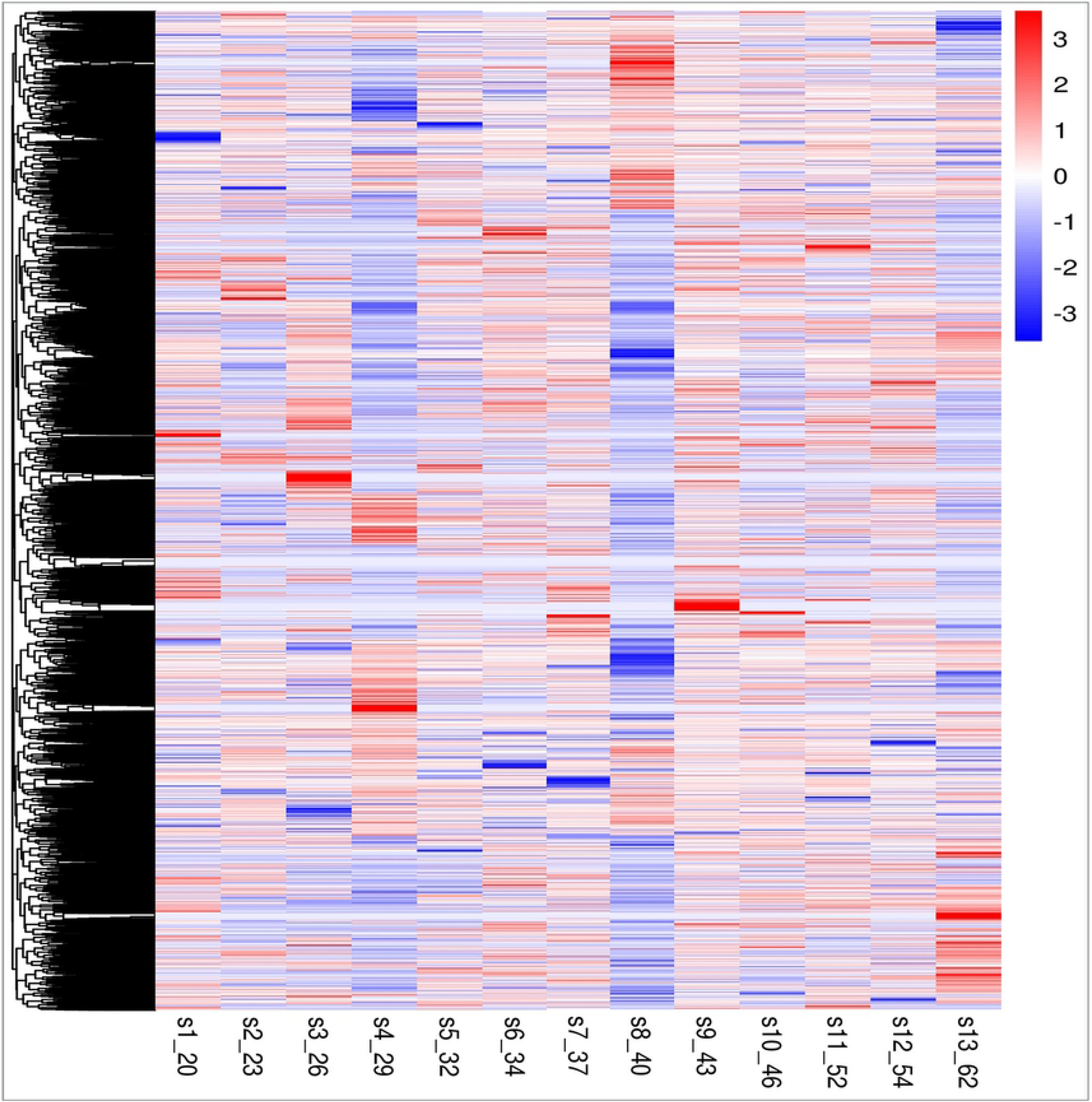
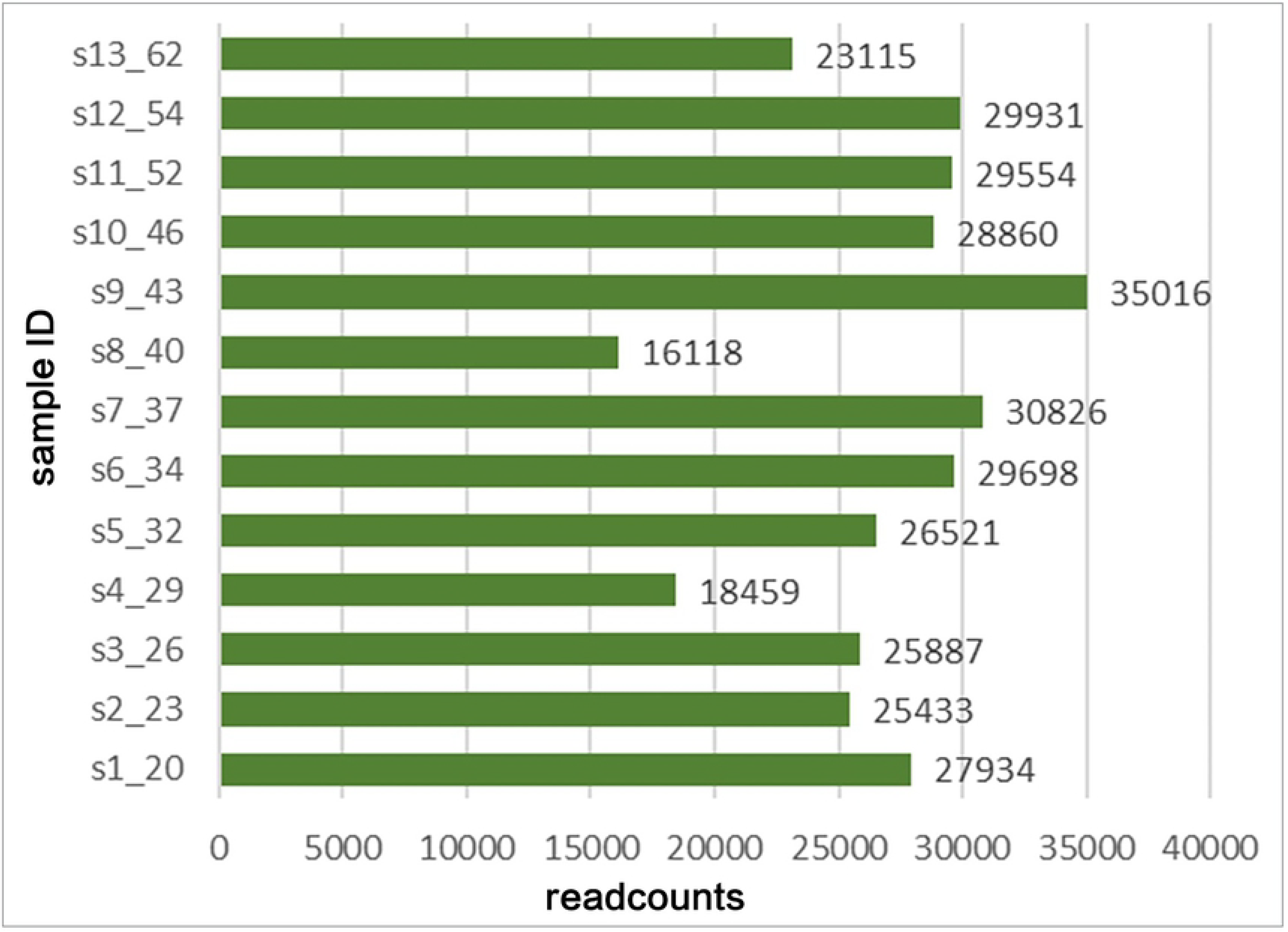
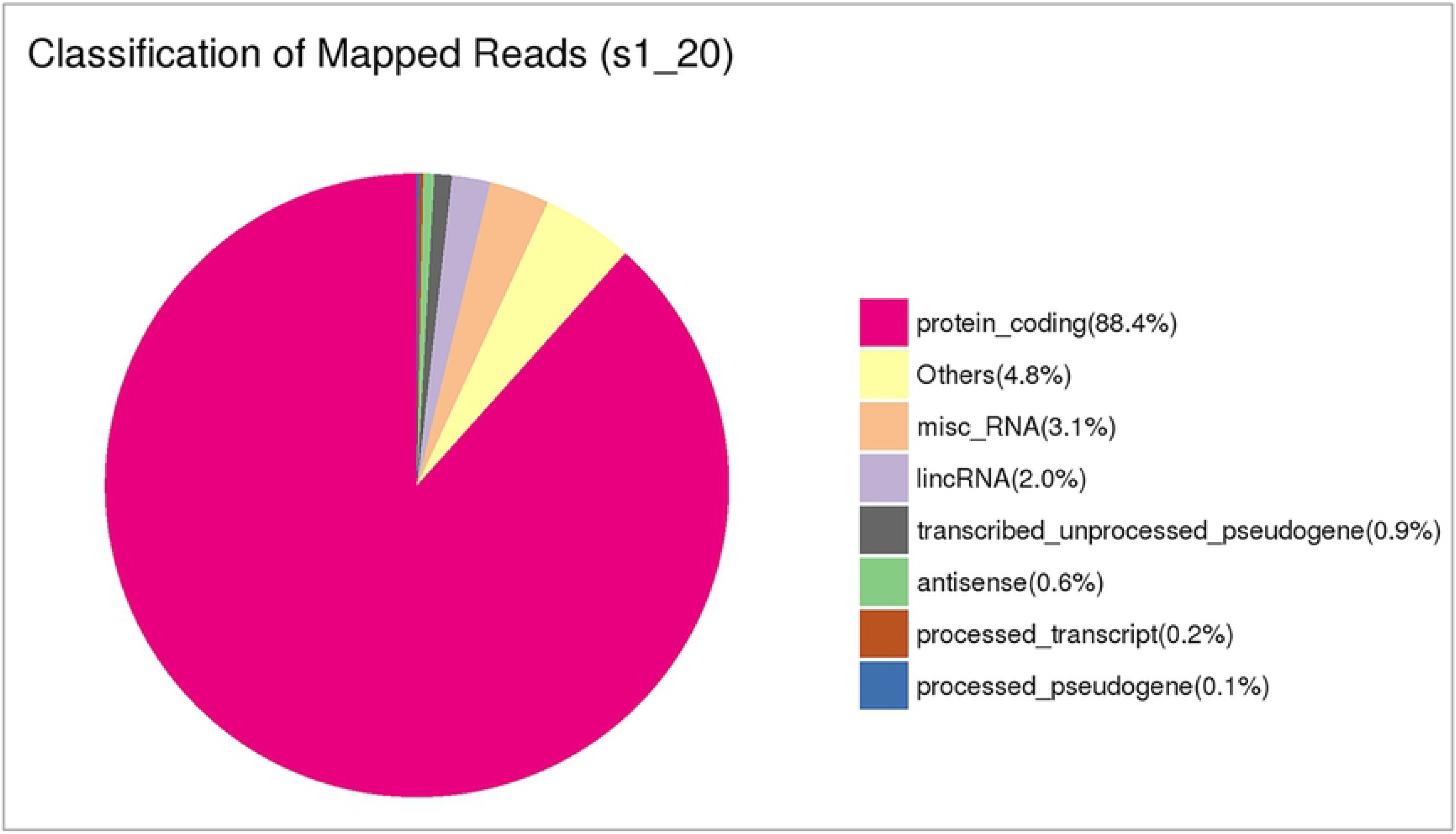
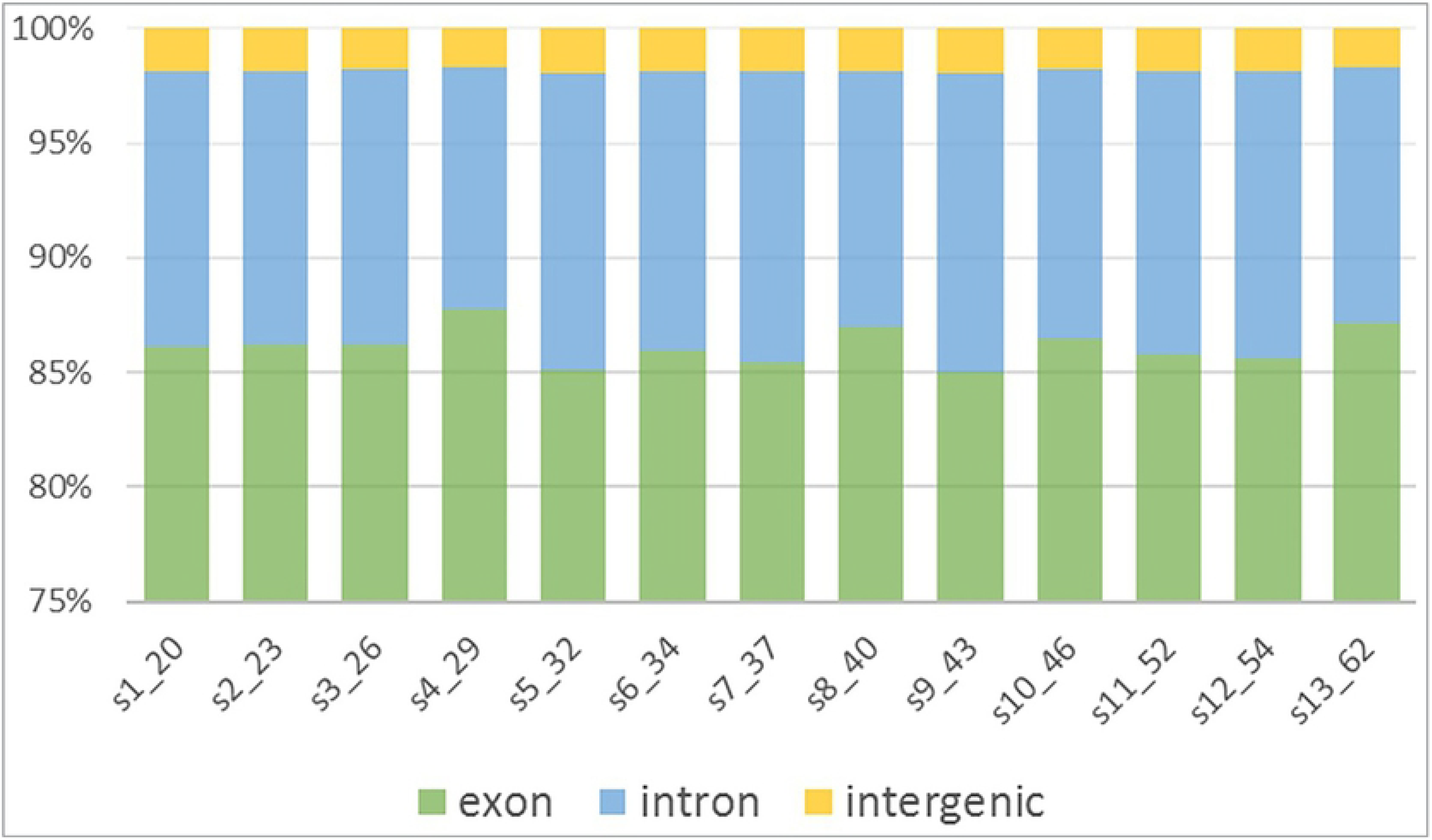
Landscape of circRNAs profile in all samples. (a) Heatmap of circRNAs expression across all 13 samples; (b) Number of circRNAs detected in all 13 samples by RNA-seq. circRNA expression analysis clearly demonstrated high abundance in human peripheral whole blood with an average number of 26719. (c) Classification of mapped reads taking sample 1 (20 years old) for example. circRNAs distributed across various genomic regions, but most commonly from protein coding regions, where over 80% of circRNAs originated. (d) Genomic features of circRNAs in each sample. Exons and introns account for higher proportion (more than 90%) than intergenic regions.

To identify age-correlated circRNA candidates, we introduced three methods of feature dimension reduction, including false discovery rate (FDR), lasso regression (LASSO), and support vector machine (SVM) (Fig 2). Spearman’s correlation coefficient (R) was calculated to identify the correlation between age and circRNA expression level (TPM value) for each single circRNA. For bivariate linear correlation analysis, 14 circRNAs were considered as age-related markers as they satisfied the following criteria: absolute R-value ≥ 0.6 and false discovery rate-adjusted p-value ≤ 0.01. Six circRNAs were selected using lasso regression method from 197 circRNAs with an absolute correlation coefficient ≥ 0.6 (S3 Table). SVM method selected 9 circRNAs from more than 40,000 circRNAs. A subset of these 28 circRNAs was chosen for age-associated circRNA candidates for further RT-qPCR validation (S4 Table).

**Fig 2.**
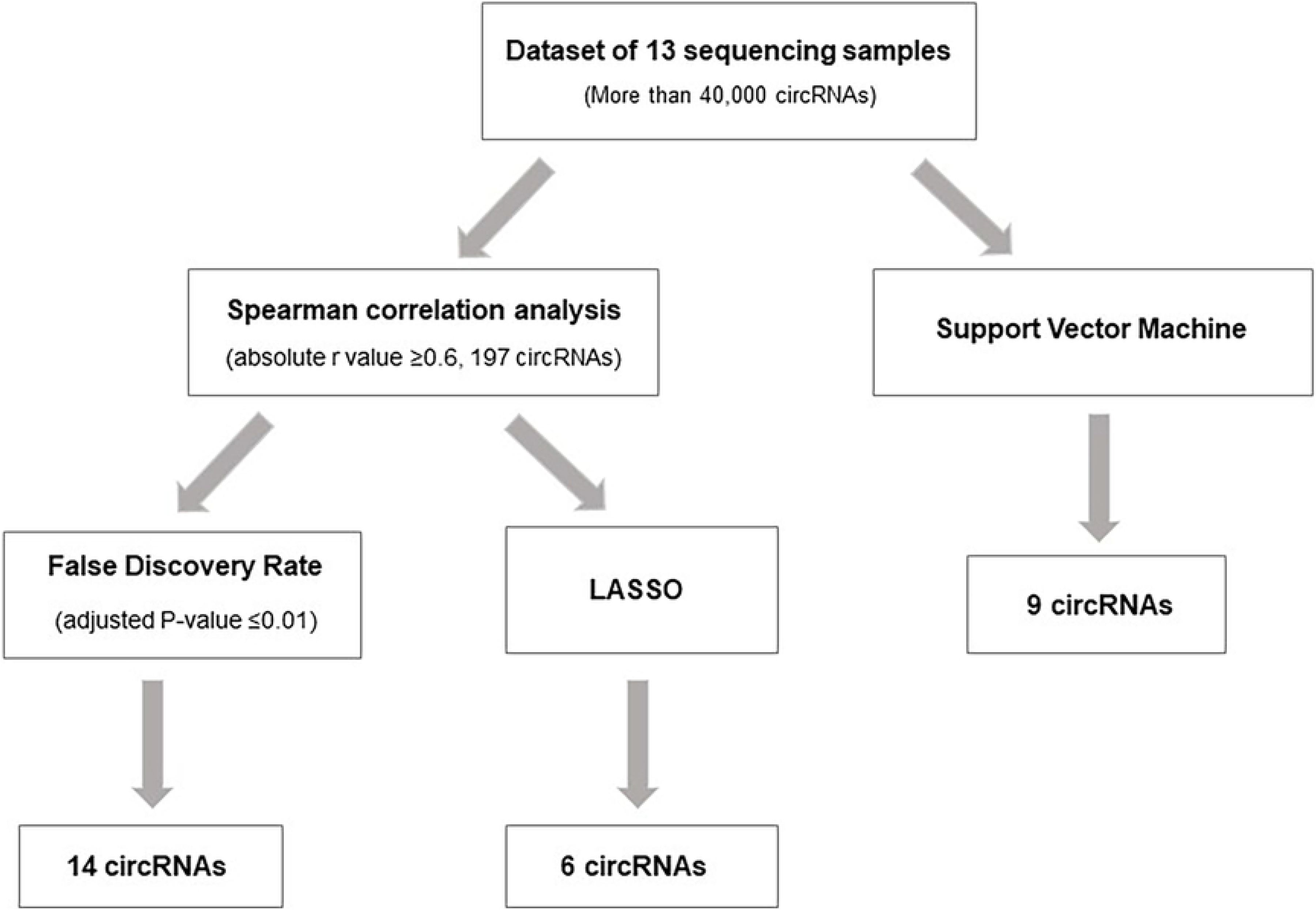
Workflow of age-related circRNAs selection.

### Validation of age-correlated circRNA using RT-qPCR

To further investigate whether the circRNA expression pattern was present similarly in other individuals. We detected 28 circRNA candidates of 50 biologically independent individuals aged between 19 years and 72 years (Fig 3a) using the Real-time fluorescence quantitative PCR (RT-qPCR) strategy. Circular RNAs are circular in shape and covalently closed. These unique transcripts are often generated by back-splicing events. They do not contain a 5’ or 3’ end nor do they have a poly-A tail. Therefore, primer design is of vital importance for PCR quantitation. The use of divergent, rather than convergent primers can selectively detect and quantitate these special RNA molecules (Fig 3b). In addition, to confirm a subset of age-related circRNA candidates, we treated the total RNA of them and their mRNA counterparts with the exoribonuclease RNase R which can reduce levels of mRNAs but not of circRNAs to provide evidence of circRNAs’ nonlinearity. Quantitative PCR (qPCR) tests were performed on the control group and RNase R treated group using convergent primers (amplify mRNA species) and divergent primers (amplify circular RNA species). It turned out that linear species and circular species were tend to be decreased by RNase R to a certain extent. Nevertheless, most circular RNAs were affected more mildly than their linear counterparts which were substantially degraded by RNase R (Fig 3c).

**Fig 3.**
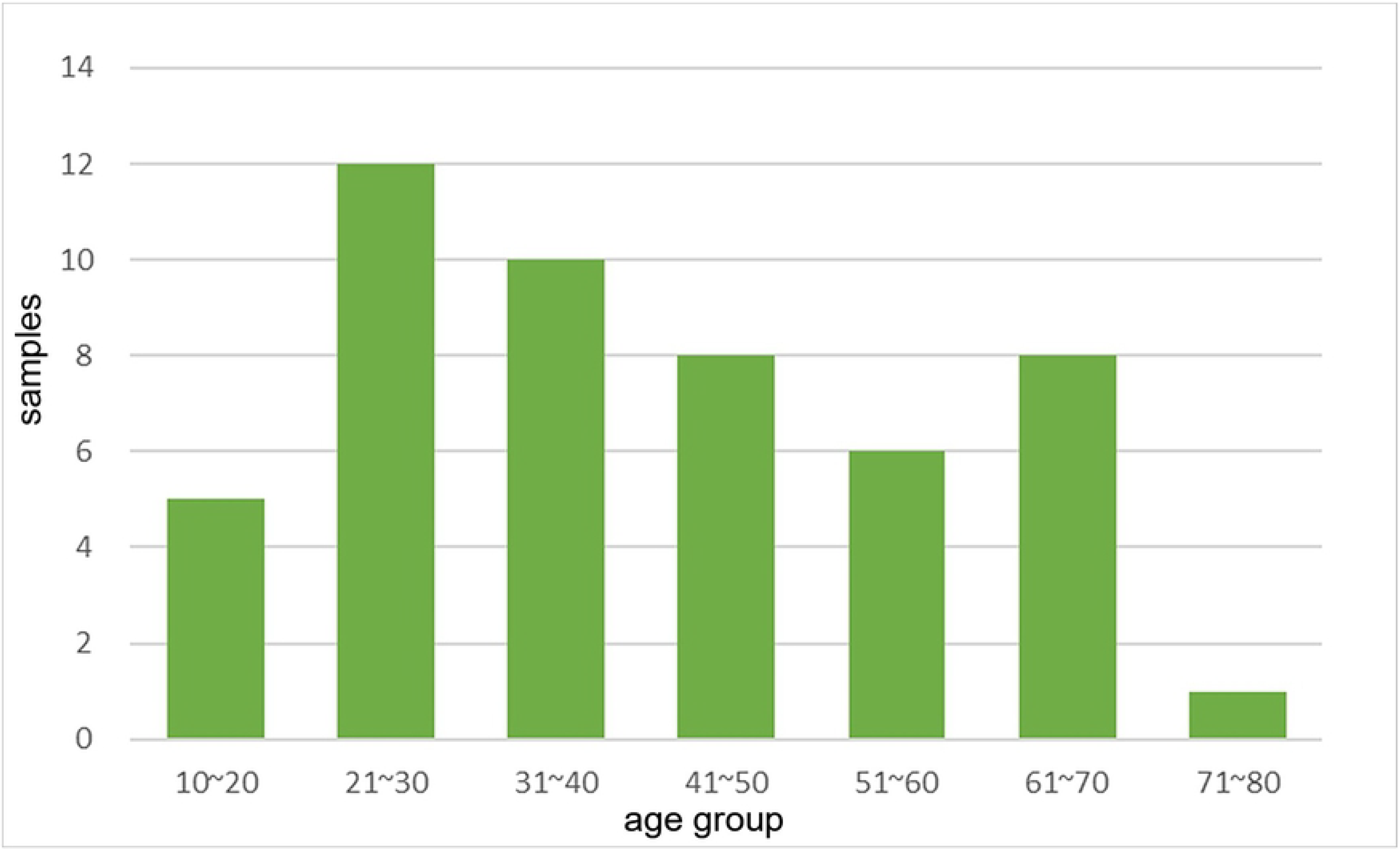
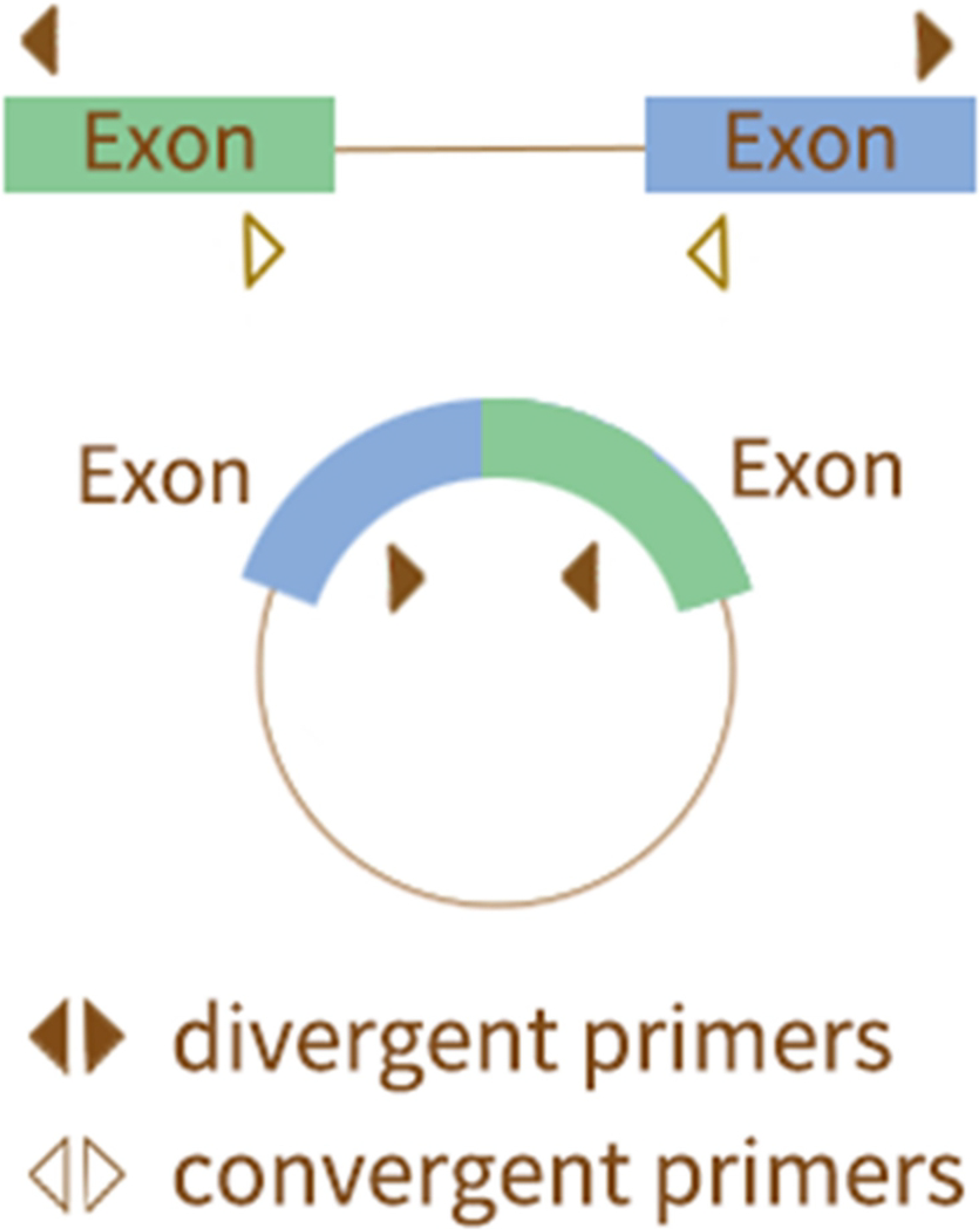
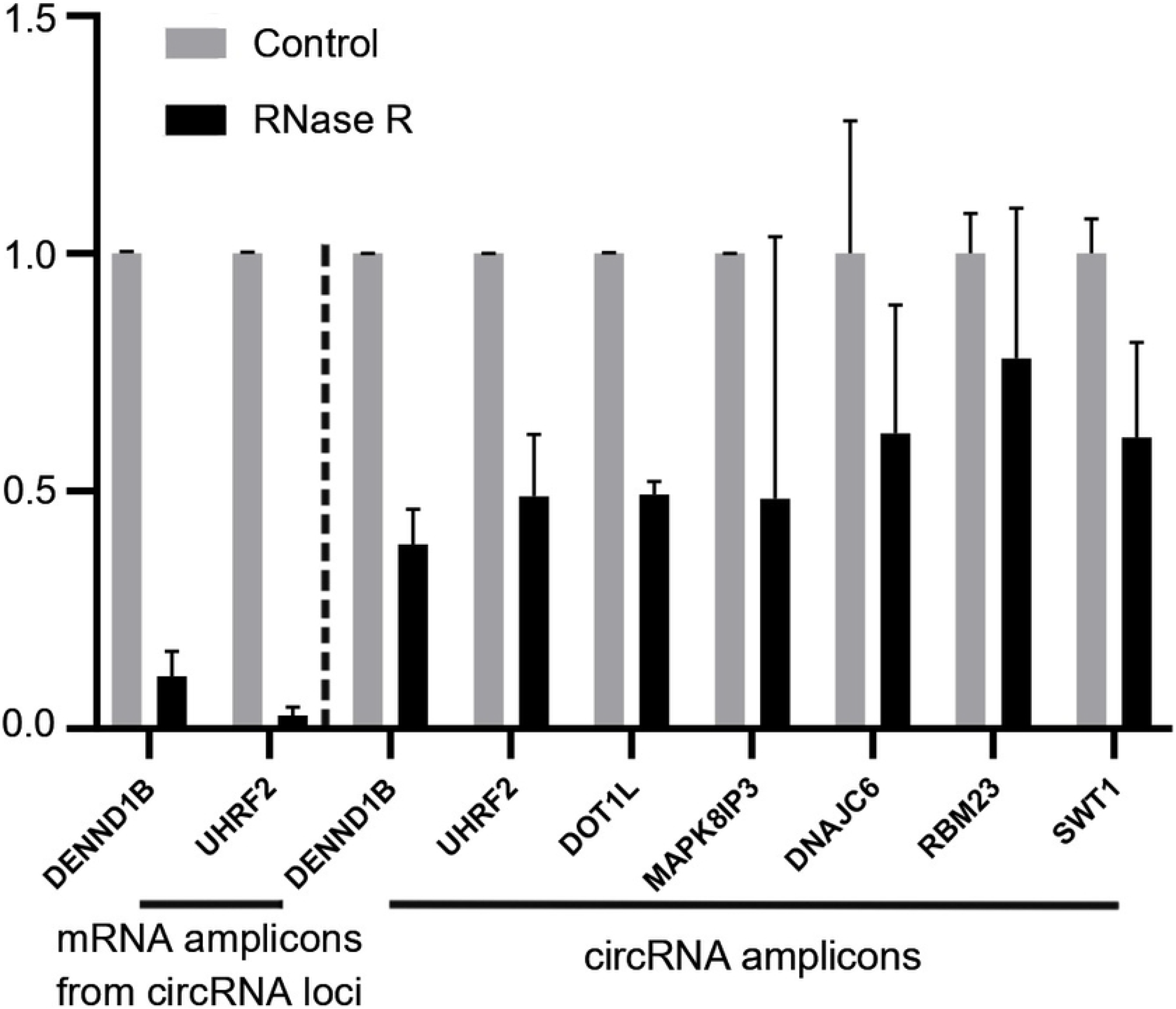
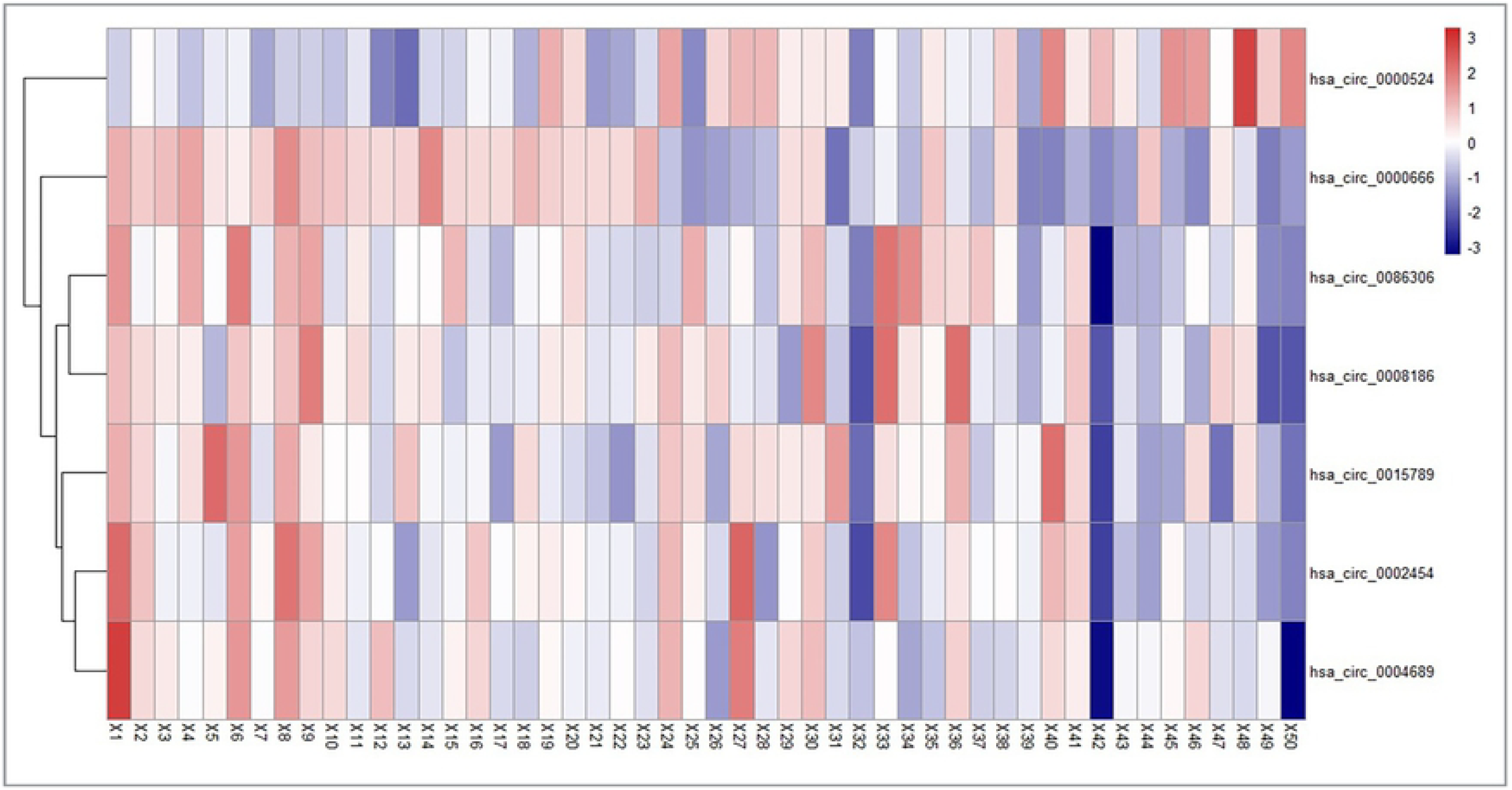
Sample distribution, principle of circRNA primer design and heatmap of age-related circRNAs. (a) Histogram of the age distribution for 50 healthy volunteers. The x-axis represents the chronological age of the individuals (age unit is years) and the y-axis (counts) represents the number of individuals. (b) Principle of circRNA primer design. The use of divergent, rather than convergent primers can selectively detect and quantitate these special RNA molecules. (c) qPCR analysis of RNase R treated samples and control samples. Most circular RNA amplicons were degraded milder than mRNA amplicons. (d) Heatmap of circRNAs expression and hierarchical clustering of 50 samples for 7 age-associated circRNA candidates from RT-qPCR. 7 out of 28 circRNA candidates change with ageing among 50 healthy unrelated individuals. All these 7 circRNAs change positively with age except for hsa_circ_0000524.

Experimentally validated circRNAs were calculated Spearman’s correlation coefficients to select age-related circRNA. Only 7 circRNAs detected by RNA-seq were in good agreement with RT-qPCR quantifications (Fig 3d). An overwhelming bias for the upregulation of circRNAs during aging was uncovered that these 7 circRNAs showing a statistically different change by qPCR were upregulated during aging except hsa_circ_0000524. The Spearman correlation coefficients among 7 circRNAs were presented in the scatter plots (Fig 4). Among them, 2 circRNAs showed slightly higher Spearman correlation coefficients (hsa_circ_0000666: R = −0.704, hsa_circ_0000666: R = 0.527). The 7 age-related circRNAs were derived from seven different genes (*DENND1B, UHRF2, DOT1L, MAPK8IP3, DNAJC6, RBM23* and *SWT1*), encoding several proteins or involving in multiple pathways. The feature description of 7 age-associated circRNAs was listed in Table 1. Furthermore, this expression trend was in accordance with previous research reported in other organisms that most circRNA levels dramatically increase during aging[32]. Finally, 7 age-correlated circRNAs were regarded as candidates for further age estimation modeling.

**Table 1.** Description of circRNA markers in the final age-prediction models.

| circRNA ID | Gene | Full gene name | Chr. | Spliced length | Change with age |
| --- | --- | --- | --- | --- | --- |
| hsa_circ_0015789 | DENND1B | DENN domain containing 1B | 1 | 477 nt | up |
| hsa_circ_0086306 | UHRF2 | Ubiquitin like with PHD and ring finger domains 2 | 9 | 297 nt | up |
| hsa_circ_0008186 | DOT1L | DOT1 like histone lysine methyltransferase | 19 | 412 nt | up |
| hsa_circ_0000666 | MAPK8IP3 | Mitogen-activated protein kinase 8 interacting protein 3 | 16 | 469 nt | up |
| hsa_circ_0002454 | DNAJC6 | DnaJ heat shock protein family (Hsp40) member C6 | 1 | 350 nt | up |
| hsa_circ_0000524 | RBM23 | RNA binding motif protein 23 | 14 | 189 nt | down |
| hsa_circ_0004689 | SWT1 | SWT1 RNA endoribonuclease homolog | 1 | 469 nt | up |

**Fig 4.**
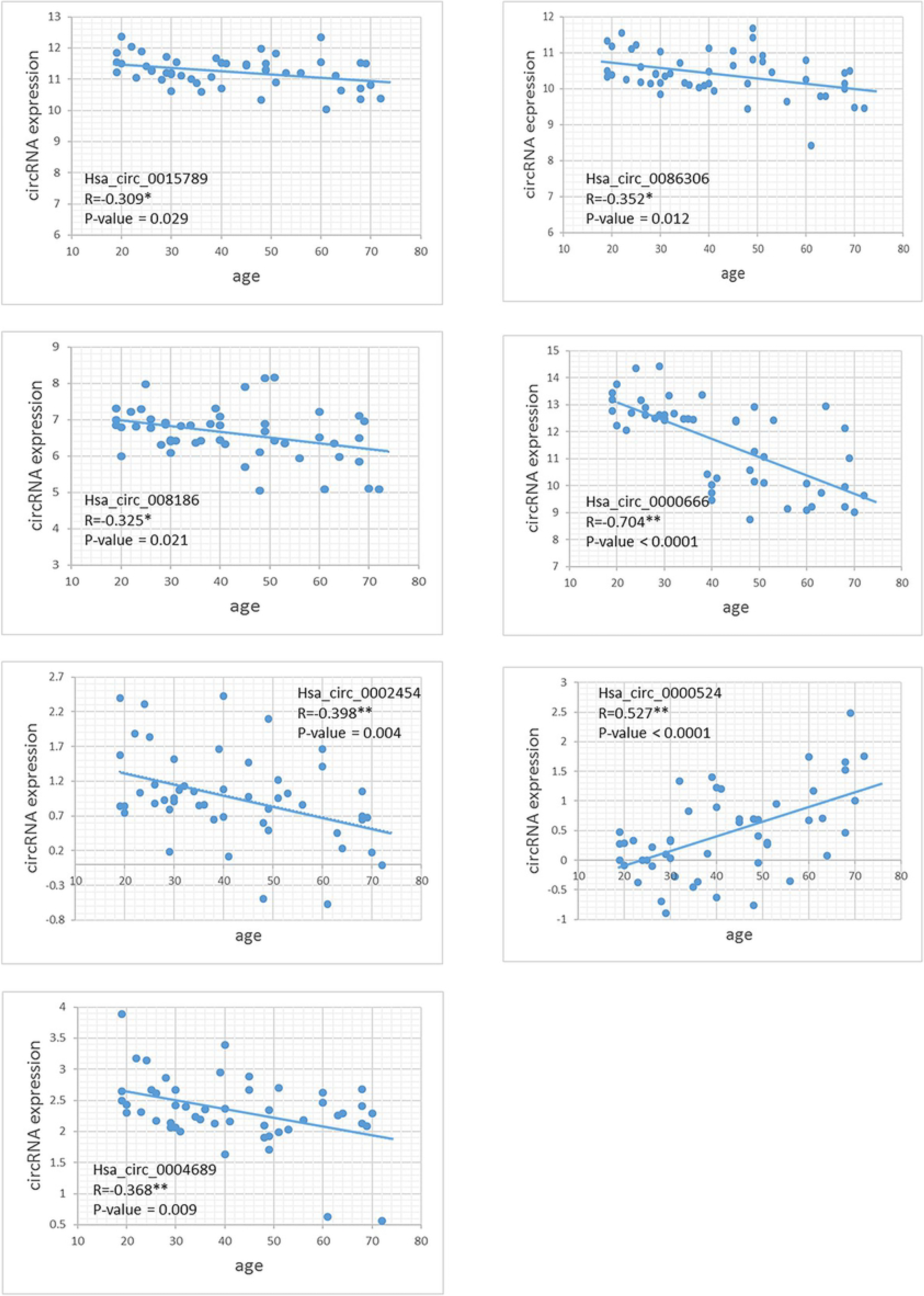
Scatter plots of ΔCt-value versus age for 7 age-related circRNAs. Scatter plots of ΔCt-value versus age for 7 age-related circRNAs. Among these circRNAs, six are positive associated with age and only one is negatively associated with age. One of them achieves the highest R-value (hsa_circ_0000666, rho = −0.704***, P<0.000).

### Development of age prediction models using different algorithms

First of all, multivariate analysis was tested to assess the effect of gender on age-associated circRNAs, no significant correlation was determined (p = 0.80). To provide an unbiased estimate of predictive accuracy for age, various attempts had been made to develop better models to predict age. We adopted five different algorithms, including multiple linear regression (MLR), bagging regression, regression tree, random forest regression (RFR), support vector regression (SVR). We fitted age prediction models based on 7 age-related circRNAs from the qPCR experiment dataset of 50 samples. Results uncovered that the regression tree model showed the lowest accuracy with a MAE of 8.474 years (R^2^ = 0.582), while RFR reached the highest accuracy (MAE = 4.096, R^2^ = 0.900). Other models showed medium MAE values, 4.155 years for SVM, 6.874 years for bagging, and 7.788 years for MLR (Table 2). Fig 5 shows the relationship between the chronological and the predicted age using different algorithms. Real and predicted ages were significantly related with a correlation coefficient value (R) of 0.953 for the RFR model (Fig 5b), indicating that the RFR algorithm seems to use weak markers to create a stronger model. The RFR model fitted 68% (34/50) of individuals within a ±5 years error range, while 94% (47/50) within a ±10 years error range. Additionally, the variable importance measures (VIM) ranking the variables (i.e. the features) with respect to their relevance for prediction is a byproduct of random forest. VIM is one of methods in capturing the patterns of dependency between variables and response in the form of a single number. It can essentially be used to many prediction methods but particularly effective for black-box methods which (in contrast to, say, generalized linear models) yield less interpretable results[33]. As an illustration, the rank of variable importance measures according to accuracy, MSE and standard errors were displayed in Fig 6, separately. Two important parameters: *mtry* referring to the number of features to be considered at each split and *ntree* standing for the number of trees in a forest were set to 3 and 500, respectively.

**Table 2.** Comparison of five regression models.

| Model | MAE<br>(years) | RMSE<br>(years) | $R^2$ | $\pm 5$ years | | $\pm 10$ years | |
| --- | --- | --- | --- | --- | --- | --- | --- |
|  |  |  |  | n | % | n | % |
| Multiple linear regression | 7.788 | 9.249 | 0.669 | 17 | 34% | 36 | 72% |
| Regression tree | 8.474 | 10.384 | 0.582 | 16 | 32% | 35 | 70% |
| Bagging regression | 6.874 | 8.839 | 0.697 | 22 | 44% | 36 | 72% |
| Random forest regression | 4.096 | 5.093 | 0.900 | 34 | 68% | 47 | 94% |
| Support vector regression | 4.155 | 6.015 | 0.860 | 39 | 78% | 45 | 90% |

**Fig 5.**
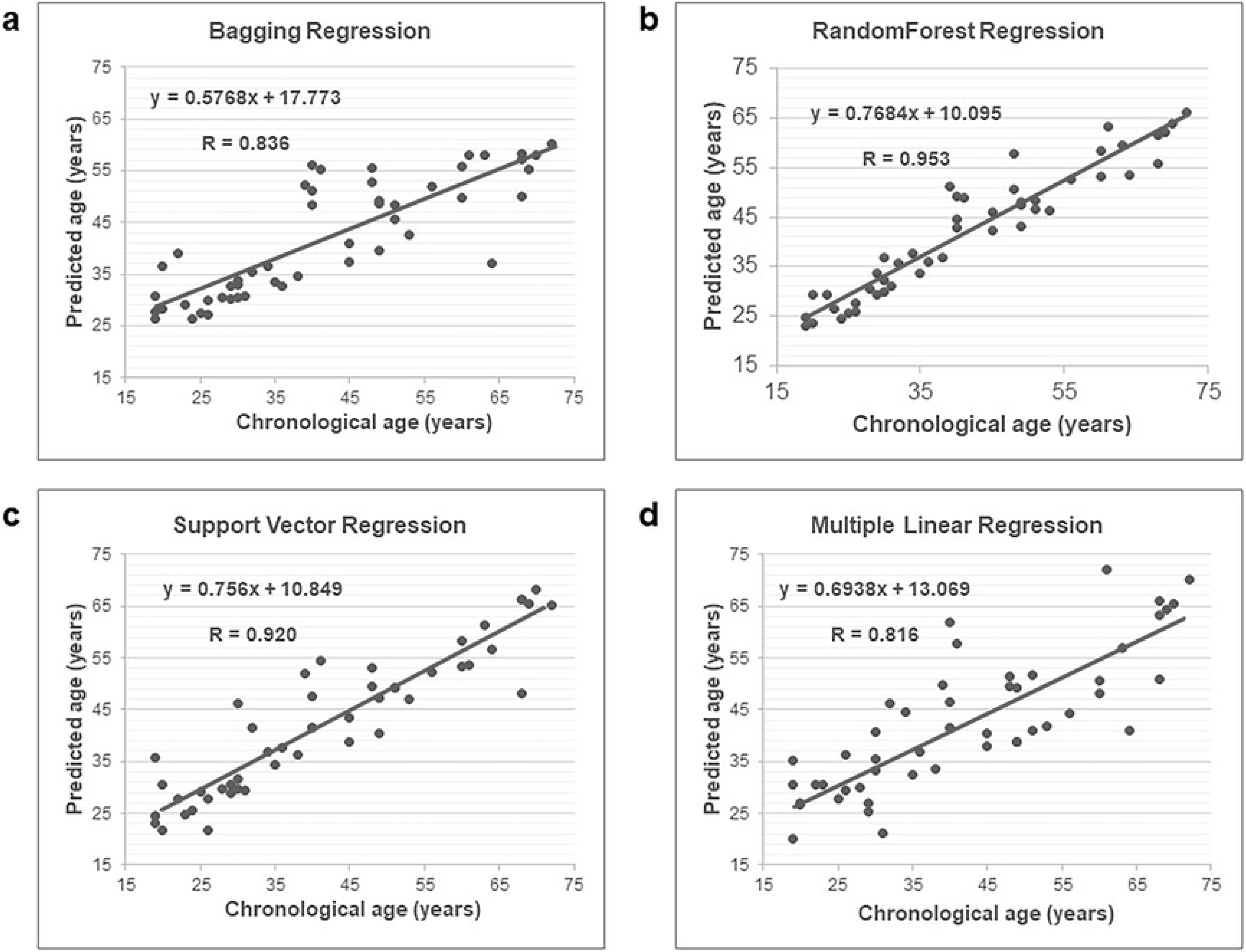
Scatter plots between chronological versus predict age using different machine learning algorithms. Predicted versus chronological ages using (a) bagging regression, (b) random forest regression (c) support vector regression (d) multiple linear regression. RFR model showed the highest connection between real and predicted age with a correlation coefficient value (R) of 0.953.

**Fig 6.**
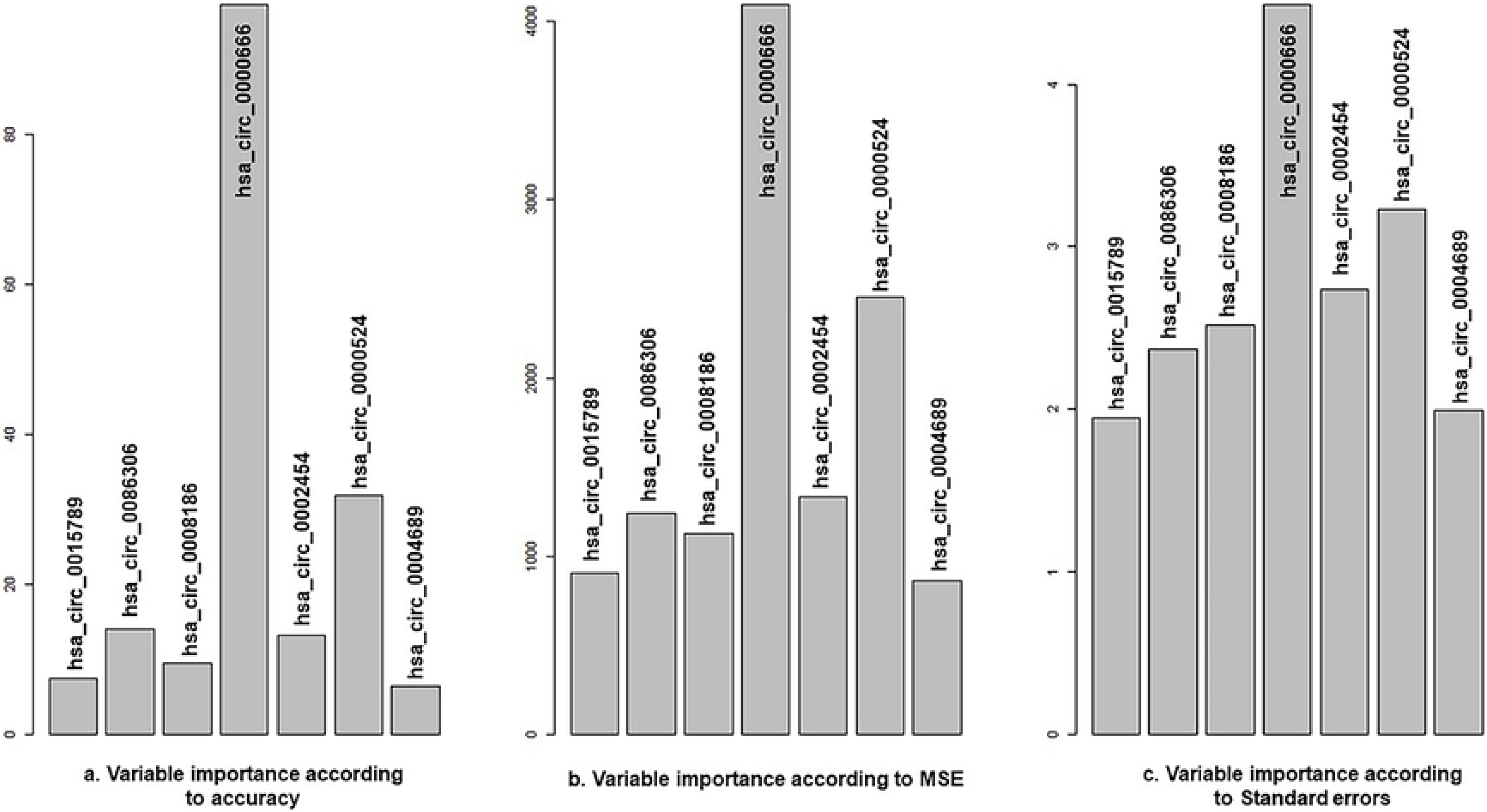
Variable importance measures (VIM) of RFR model. Variable importance measures (VIM) of RFR model according to (a) accuracy, (b) MSE and (c) standard errors. circRNA hsa_circ_0000666 is the most important feature among all seven features, followed by hsa_circ_0000524.

In order to further evaluate the predictive error, samples were divided into five age groups: 19-30 years, 31-40 years, 41-50 years, 51-60 years and 61-72 years. In line with previous studies, MAE increased with age[30]. In our study, older individuals (>60 years old) presented an increased deviation between chronological and predicted age compared to younger groups (<40 years old) and the predicted age of elders was prone to be underestimated. Specifically, the RFR method, for example, MAE for the youngest age group (19-30 years old) was 3.479 years and 6.657 years for the oldest (61-72 years old) (Table 3).

**Table 3.** MAE for different age categories (RFR model)

| Age ranges | N | MAE |
| --- | --- | --- |
| 19-30 years | 17 | 3.479 years |
| 31-40 years | 10 | 3.895 years |
| 41-50 years | 8 | 4.049 years |
| 51-60 years | 6 | 4.267 years |
| 61-72 years | 9 | 6.657 years |

A 10-fold cross validation was performed to further test the accuracy of the models. In a 10-fold cross-validation (CV), the original dataset is randomly separated into 10 subsets of approximately equal sizes. After each of the iterations, one of the folds (5 samples in the current study) was chosen as the validation set, while the remaining 45 samples were used for training. Results showed that MAE changed slightly after a 10-fold CV but very close to the MAE from the whole dataset. MAE was decreased to 4.064 years for RFR (R^2^ = 0.902), 4.155 years for SVR (R^2^ = 0.867), 7.634 years for MLR (R^2^ = 0.664), and increased to 9.401 years for regression tree (R^2^ = 0.498) and 7.253 years for bagging regression (R^2^ = 0.670) (S5 Table).

## Discussion

Although previous studies mostly relied on morphological methods, the development of molecular methods for age prediction is extremely valuable when the human specimens retrieved from crime scenes without morphological age features, such as bloodstain which is commonly seen. Considerable progress of the molecular biology of aging has been made in the last decades and using a variety of age-associated biomarkers in blood has emerged as a useful method for molecular age estimation. Nevertheless, estimating age accurately and reliably from molecular biomarkers is still a challenge because identifying minimal numbers of biomarkers that provide maximal age informative is extremely tough when analyzing forensic samples as the nature of forensic DNA or RNA being of lower quality and quantity[2]. New insights into discovering novel biomarkers with property of strong correlation of aging and stability need more attention within the forensic genetic community.

In this preliminary study, we provided potential age-associated markers in human blood, namely circRNA due to its property of highly stable and developmental stage-specific expression patterns. To the best of our knowledge, this is the first one to predict an individual’s age using circRNA biomarkers. CircRNAs originate through back-splicing events from linear primary transcripts, presenting as a special covalent loop without a 5’ cap or 3’ tail. Recently some analyses conducted by forensic researchers have focused on the capacity of circRNAs to identify body fluid due to their tissue-specificity[34]. However, we noticed that circRNAs have been studied in aging and age-related diseases across different species, revealing a global accumulation of circRNAs during aging. Thus, we came up with a hypothesis if similar age-related circRNAs accumulation is also observable in human blood. Additionally, the unique feature of circRNAs enhances their stability thus might be more suitable for forensic degraded samples seen commonly. To substantiate this assumption, in the current study, we identified age-dependent circRNA markers from human peripheral whole blood by means of circRNA next-generation sequencing and real-time PCR and developed age-predictive models using different machine learning algorithms. Our study is none-trivial, we opened a new avenue for forensic age prediction analysis using a novel biomarker in human blood, which would be an important topic for future investigation.

Selecting truly age-associated circRNAs from a giant gene dataset of 13 individuals’ circRNA-seq profiles (approximately over 40,000 circRNAs were identified) made the current study difficult. Recent years, sequencing technological advances in monitoring gene expression have given rise to a dramatic increase in gene expression data and machine learning techniques are known to be excellent at discarding and compacting redundant information from gene expression data. The introduction of machine learning algorithms helped us to select variables from 45,697 down to 28 effectively. 7 out of 28 age-dependent circRNAs were finally selected according to Spearman’s correlation coefficients calculation and p-value. They showed positive correlations with chronological age except for hsa_circ_0000666. This expression trend was in accordance with previous research in other organisms that most circRNA levels dramatically increase during aging[35]. Though there has no existing evidence yet indicating that circRNA expression levels of human peripheral whole blood are changing with increasing age, recent reports showed that circRNAs increase in the elders’ brains and play a vital role in the development of neurodegenerative diseases like Alzheimer’s disease (AD) and Parkinson’s disease (PD)[36–38]. Furthermore, Shahnaz Haque et al. assessed circRNA expression in aging human blood and found that circFOXO3 and circEP300 demonstrated differential expression in one or more human senescent cell types, providing indirect evidence for circRNA as a promising indicator for aging[26].

Selected circRNA candidates in our study were derived from seven different genes (*DENND1B, UHRF2, DOT1L, MAPK8IP3, DNAJC6, RBM23* and *SWT1*) and might involve in several important the biological processes, we then undertook a comprehensive analysis. As expected, these seven genes code for important proteins or involve in biological metabolic process. Analyses were performed on string-db.org and the gene database of www.ncbi.nlm.nih.gov. The gene *DENND1B* encodes guanine nucleotide exchange factor (GEF) for RAB35 that acts as a regulator of T-cell receptor (TCR) internalization in TH2 cells which functions as a regulator in the process of childhood asthma[39]. The gene *UHRF2* encodes a protein that is an intermolecular hub protein in the cell cycle network[40]. Through cooperative DNA and histone binding, may contribute to tighter epigenetic control of gene expression in differentiated cells. Recent research found that *UHRF2* can promote DNA damage response[41]. The gene *DOT1L*, the protein encoded by this gene is a histone methyltransferase that shows significant histone methyltransferase activity against nucleosomes[42]. The gene *MAPK8IP3* encodes a regulator of the pancreatic beta-cell function. This gene is found to be mutated in a type 2 diabetes family and thus is thought to be a susceptibility gene for type 2 diabetes[43]. The gene *DNAJC6* belongs to the evolutionarily conserved DNAJ/HSP40 family of proteins, which regulate molecular chaperone activity by stimulating ATPase activity. This gene plays a role in clathrin-mediated endocytosis in neurons[44]. The gene *RBM23* encodes probable RNA-binding protein and may be involved in the pre-mRNA splicing process[45]. *SWT1* is a protein coding gene that involves in diseases including kidney sarcoma and Wilms tumor 1. Notably, the function of circRNAs or their host genes can provide some clues for future selection of biomarker candidates. Some circRNAs were confirmed to regulate biogenesis and some senescence-associated genes might become a clue for the selection of age-dependent indicators, for instance, gene *Foxo3* circular RNA was highly expressed in heart samples of aged patients and mice reported by William W. Du et al[46]. and was found different expression in one or more human senescent cell types in the study of Shahnaz Haque et al.[26]. In contrast, some of the aforementioned circRNAs play a key role in the pathogenesis of tumors or diseases such as *DENND1B, MAPK8IP3* and *SWT1*. Hence, the effect of disease state on age predictions should be considered for further investigation because it is important to build a robust predictive model using biomarkers that would not present differentially expression due to disease status. As models described in Athina Vidaki et al.[1], they analyzed the effect of different diseases on age prediction that schizophrenia presented the lowest age prediction error, while anemia demonstrated the lower relation with age, indicating that it becomes evident that the error is much higher for blood related diseases when analyzing separately samples suffering from blood *vs.* non-blood related diseases.

Among the multiple machine learning algorithms adopted for the present study, RFR outperformed the others with a RMSE value of 5.072 years and a MAE value of 4.065 years (R^2^ = 0.902), followed by SVR with a MAE of 4.166 years and a RMSE of 5.936 years (R^2^ = 0.865). This estimation errors were close to previous predictive models using other biomarkers[8, 15]. RFR model explained 94% of the variation in real and predicted age (±10 years) with the accuracy of a MAE of 3.479 years in the ≤30 years age group and 6.657 years in the older group (≥ 61 years). Our models perform better in young subjects but poorly for elders over 60 years old. We suppose that the deviation might attribute to the fact that adults and elders suffer from more confounding factors in the aspect of medication, smoking, or alcohol, which are not easily accessible for children and adolescents, as previously reported[14]. Another possible reason might be the smaller sample size included in our study especially elder samples, which leads to bias. We are aware of one of the weaknesses in the current study that our sample set did not incorporate children and adolescents (a population sample ranging from 19-72 years old was used). Therefore, future studies are needed to explore age-specific biomarkers, namely under-aged-specific and adult-specific predictive indicators. Focusing on when these biomarkers occur and when they lose functions or their periodical changes during an individual’s life span also need to be investigated precisely in further research.

Another limitation of the current study is a pretty limited sample size, which would restrict our models’ validity to a certain extent. Although machine learning algorithms seem to achieve a relatively high accuracy using a subset of weak predictors. The prediction accuracy of models presented here is needed to be further improved to satisfy forensic practice, which means more robust, stable and higher age-related circRNA biomarkers need to be found.

It is worth asking what underlying mechanisms might give rise to the increased levels of circRNAs with aging? Some people hold that the age-related increase tends for circRNAs are reflective of age-accumulation more so than specific regulation. Such an assumption is based on studies of multiple animals discovering that the increased abundant of circRNAs during aging was found to be largely independent of gene expression from their host due to their lack of 3’ or 5’ end[32]. In addition to the contribution of circRNA stability, some researchers suppose that increased circRNA biogenesis due to age-related changes in alternative splicing might play a role[47]. These results prompt us that it might be necessary to figure out the functions of circRNAs in aging and mechanisms contributing to circRNA changes during aging in future work.

Our study though suffers from some drawbacks, the proposed age prediction models here exhibit good estimation accuracy and provide strong evidence for circRNAs as potential biomarkers to be applied in practice in trap future. Nevertheless, it is convinced that prediction can be improved in our future work through optimized algorithms, enlarged sample size and a wide age range included. In a word, based on our results, we argue that the prediction of the chronological age utilizing age-dependent changes of specific circRNAs is a promising application and will become an increasing field of interest.

## Materials and methods

### Samples collection and RNA isolation

Whole blood samples were collected from 13 healthy volunteers (aged between 20 and 62 years) (S2 Table) for circRNA sequencing and 50 (aged between 19 and 72 years) for RT-qPCR experiments validation. Written informed consent was obtained from all the volunteers, and our study was approved by the Medical Ethics Committee of Hebei Medical University (No. 20190013).

Peripheral whole blood 10ml were drawn from subjects by venipuncture and collected in an EDTA-containing vacutainer. Total RNA was isolated immediately using TRIzol reagent (Thermo Scientific, USA) according to the manufacturer’s instructions after blood collection. RNA quantification was conducted on the NanoDrop 1000 (NanoDrop Technologies). RNA integrity was assessed using the RNA Nano 6000 Assay Kit of Bioanalyzer 2100 system (Agilent Technologies, CA, USA). Isolated RNAs were preserved on the condition of −80°C until reverse transcription.

### High throughput sequencing (circRNA-seq)

A total amount of 5 μg RNA per sample was used as input material for RNA sample preparation. Firstly, ribosomal RNA was removed by Epicetre Ribozero™ rRNA Removal Kit (Epicentre, USA), and rRNA free residue was cleaned up by ethanol precipitation. Subsequently, the linear RNA was digested with 3U of RNase R (Epicentre, USA) per μg of RNA. The sequencing libraries were generated by NEBNext^®^ Ultra™ Directional RNA Library recommendations. Barcoded libraries were sequenced at Novogene Co., LTD (Beijing, China) using the Illumina Hiseq 4000 platform to obtain paired-end 150nt reads. circRNA candidates were detected and identified using find_circ algorithm[48]. The normalization of contig counts was performed by calculating transcripts per million (TPMs). The normalized expression level = (read counts *1,000,000) / libsize (libsize is the sum of circRNA read counts).

### Reverse transcription and Quantitative PCR (RT-qPCR)

RNA was reverse transcribed into cDNA using PrimeScript Reverse Transcriptase™ (Takara Bio Inc., Otsu, Shiga, Japan) according to the manufacturer’s protocol. cDNA was stored at −80°C waiting for further RT-qPCR tests. qPCR reactions were performed using QuantiNova™ SYBR^®^ Green PCR Kit (Qiagen, Germany) on a 7500 System (Applied Biosystems). Sequences of primers are available in the S1 Table. 18S rRNA was chosen as reference genes which was stably expressed in blood samples in the pre-test. Delta Ct value (Ct_target_ – Ct_reference_) represented circRNAs expression. These tests were performed using technical quadruplicates. PCR products were also electrophoresis on agarose gel and recovered to verify the specificity of primers using Sanger sequencing. Validation tests of circular RNA nonlinearity were performed using total RNA samples with or without 2.5 U/μg RNase R for 50 min at 37 °C. Equal amounts of control samples and RNase R treated RNA served as input for cDNA preparation.

### Selection of potential age-associated circRNAs

An enormous of data produced by the next generation sequencing makes a pre-selection process very important and challenging. In an attempt to limit the quantity of predictors which may save both time and cost, three statistical methods were conducted to select age-associated markers. According to circRNA-seq data of 13 subjects in age from 20 to 62 years, we used adjusted p-value lower than 0.01 (false discovery rate method) after the Spearman’s correlation analysis as a criterion to identify the association between circRNA expression levels and chronological age. Lasso regression and support vector machine (SVM) were also applied to select circRNA candidates using TPM values of each samples. The Spearman correlation coefficients were then calculated again for 50 samples according to qPCR results to choose age-dependent circRNAs for the final models.

### Statistical analysis

The correlation between age and circRNA expression levels was assessed by the Spearman’s correlation coefficients using IBM SPSS statistics software for Windows, version 21.0. Additionally, R project for statistical computing software version 4.0.0 was employed to conduct five algorithms: multiple linear regression (lm), regression tree (rpart), bagging regression (ipred), random forest regression (randomForest) and support vector regression (rminer). The RMSE and MAE values between chronological and predicted ages were used as performance metrics for the final age prediction models using the five above-mentioned algorithms. Ten-fold cross-validation was performed for validation and optimization of the prediction models. The cross validation removes a part of samples from the overall dataset randomly and assigned as a validation set and the rest of the data is taken for a training set.

## Acknowledgments

The authors would like to thank voluntary participants that donated blood samples towards this project.

## Supporting information captions

**S1 Table.** Sequences of 28 age-related circRNAs for RT-qPCR verification.

**S2 Table.** Information of circRNA-seq samples.

**S3 Table.** 197 circRNAs with a |R| ≥ 0.6 after bivariate correlation analysis.

**S4 Table**. Twenty-eight circRNAs selected by three separate approaches.

*: identical circRNA selected by different methods

**S5 Table.** Evaluation of the accuracy of different models by 10-fold cross validation analysis.

